# Cell clusters are programmed towards a reductive metabolic state by adherence junctions

**DOI:** 10.1101/2025.10.02.680172

**Authors:** Uttkarsh Ayyangar, Janhavi Sathe, Shabbir Ahmad, Sunil Laxman

**Affiliations:** BRIC Institute for Stem Cell Science and Regenerative Medicine (BRIC inStem), GKVK Post Bellary Road, Bengaluru 560065; Max Planck Institute of Molecular Cell Biology and Genetics, Dresden, Germany; Regional Centre for Biotechnology, NCR Biotech Science Cluster, Faridabad, Haryana, 121 001

## Abstract

Solitary cells form stable clusters via cell-cell adhesion using adherens junctions. The role of these junctions in early cell-state changes as cells form clusters is unclear. Here, we uncover that the formation of cadherin junctions as cells cluster drives a ubiquitous metabolic reprogramming. This reprogramming enhances the pentose phosphate pathway (PPP) and NADPH production to augment a reductive state. Consequently, cell clusters stabilized by cadherin junctions have reduced intracellular reactive oxygen species (ROS), are resistant to exogenous ROS-inducing agents, and have reduced apoptotic markers. Mechanistically, this metabolic reprogramming is driven by the cadherin-dependent activation of NRF2. Blocking the cadherin junction-dependent metabolic program reverses clustered cells to resemble the solitary cell state, increasing cell death and enhancing sensitivity to exogenous ROS. These insights suggest a biochemical basis for adherens junctions mediating a reductive metabolic program as solitary cells form clusters, with implications for understanding multicellular organization and collective cell behavior.

## Introduction

Cell clustering by adhesion, the association of single cells to form multicell groups and stay together, is a defining feature of metazoan biology, and a feature of multicellular transitions (1–4). This formation of stable cell groups offers growth and survival advantages in metazoans and was positively selected during evolution (1, 2, 4). In metazoan cells, clustering augments unique functional paradigms during early embryonic development and in tissue morphogenesis, and is critical in disease conditions such as cancer (5). In early embryonic development, cell clustering facilitates cell state and fate decisions, intercellular communication, and cellular patterning (6, 7). Alternatively, in disease conditions such as cancer, cell clustering leads to improved resistance against anti-cancer therapy (8–10). As cells form clusters, the overall cell states change. Recent studies suggest a close coupling between the cell states and cellular metabolism, wherein changes in cellular function occur alongside metabolic reprogramming (11–14) However, these gross phenomena are difficult to interpret, since as cells form clusters and grow, there are multiple inter-linked, amplifying events - through contact-based inhibition of growth, changes in the extracellular matrix, nutrient exchange, and more. The early changes in cell state and metabolism, and their relationships therein, as cells transition towards stable, adhered clusters, remain largely unknown.

A primary, required step towards solitary cells forming a stable cell cluster is the formation of cell-cell adherens junctions (15). The formation of adherens junctions is facilitated by homotypic attachment between cell adhesion proteins - the cadherins (15, 16). Cell adhesion molecules like cadherins are ubiquitous in metazoans, and their absence or mutations lead to drastic disease conditions due to the severe adhesion defects. These junctions not only serve as attachment sites but also change cellular signalling (15, 17) thereby regulating the physiological outcomes in cells. The formation of adherens junctions in metazoan cells facilitates the formation of cell clusters, and precedes the formation of cell-ECM junctions (or focal adhesions) in early evolution and mammalian development (18, 19). However, the earliest cell state and metabolic changes that are driven by the formation of adherence junctions in cell clusters remain unclear.

In this study, we investigated early changes in cell and metabolic states as solitary cells start transitioning to form stable clusters. We find that as solitary cells cluster and form stable cadherin junctions, they reprogram their metabolism towards increased pentose phosphate pathway (PPP) flux, the production of reducing equivalents (NADPH), and an overall reductive state. This results in reduced intracellular reactive oxygen species (ROS), lower oxidative stress, and increased resistance to exogenous ROS in clustered cells. This metabolic reprogramming towards increased PPP flux and a reductive state is abolished in cell clusters by disrupting the cadherin junction. Mechanistically, this occurs through the cadherin junction-dependent activation of the NRF2 pathway, which drives this metabolic program. Thus, our data reveal that the formation of adherens junctions as solitary cells form stable clusters metabolically reprograms cells towards a reductive state. This enables cells in clusters to regulate the intracellular redox state and enhances their survival. Our data establishes a pivotal role for adherens junction formation in determining cell state transitions through metabolic control, as individual cells transition to form stable multicell groups.

## Results

### Cell clustering drives a reductive metabolic program with enhanced PPP flux and altered cell states

As cells transition from solitary individuals to cell clusters, the initial changes in cellular states and metabolism that occur in the first few hours remain unknown (Fig. 1A). We wanted to understand and address the early cellular events that occur as single cells transition to form cell clusters. To address this, we devised a clear, first principles approach to minimize confounding events. We cultured mammalian cells in monolayers, on non-TC treated culture dishes coated with poly-L-lysine (PLL), with variable dilutions of seeding in to control cell density and in turn, cell clustering (Fig. 1B). Attachment of cultured cells on PLL is based on electrostatic interactions, and therefore minimizes the complex signalling circuitry augmented by cell-ECM attachments due to engagement of integrin proteins (20, 21). To facilitate cell clustering, we first seeded U2OS (human osteosarcoma cell line) cells in high-density (∼10^5^ cells), or low-density (∼10^4^ cells). The seeding the cells in high density consistently led to clustering of the cells, while the low density seeding ensured most cells were solitary (Fig. S1A). As an additional control, we confirmed that there were no obvious changes in cell size as approximated by measuring nuclear area, a standard proxy for cell area (Fig. S1B). To avoid any potential effects of nutrient limitation in higher density conditions, the culture medium was replaced with fresh medium an hour prior to cell processing for experimentation. Importantly, to capture early events, all cells were processed within 6 hours after plating. To these cells, we compared the metabolic and cell states between solitary cells - seeded in low density, or clustered cells – seeded at high density (Fig. 1B). This experimental design ensured minimal confounding effects of nutrient availability and ECM-based junctions on the solitary and clustered cells, and ensured that the characterization was of early cell and metabolic states in these two groups (Fig. 1B).

**Figure 1:**
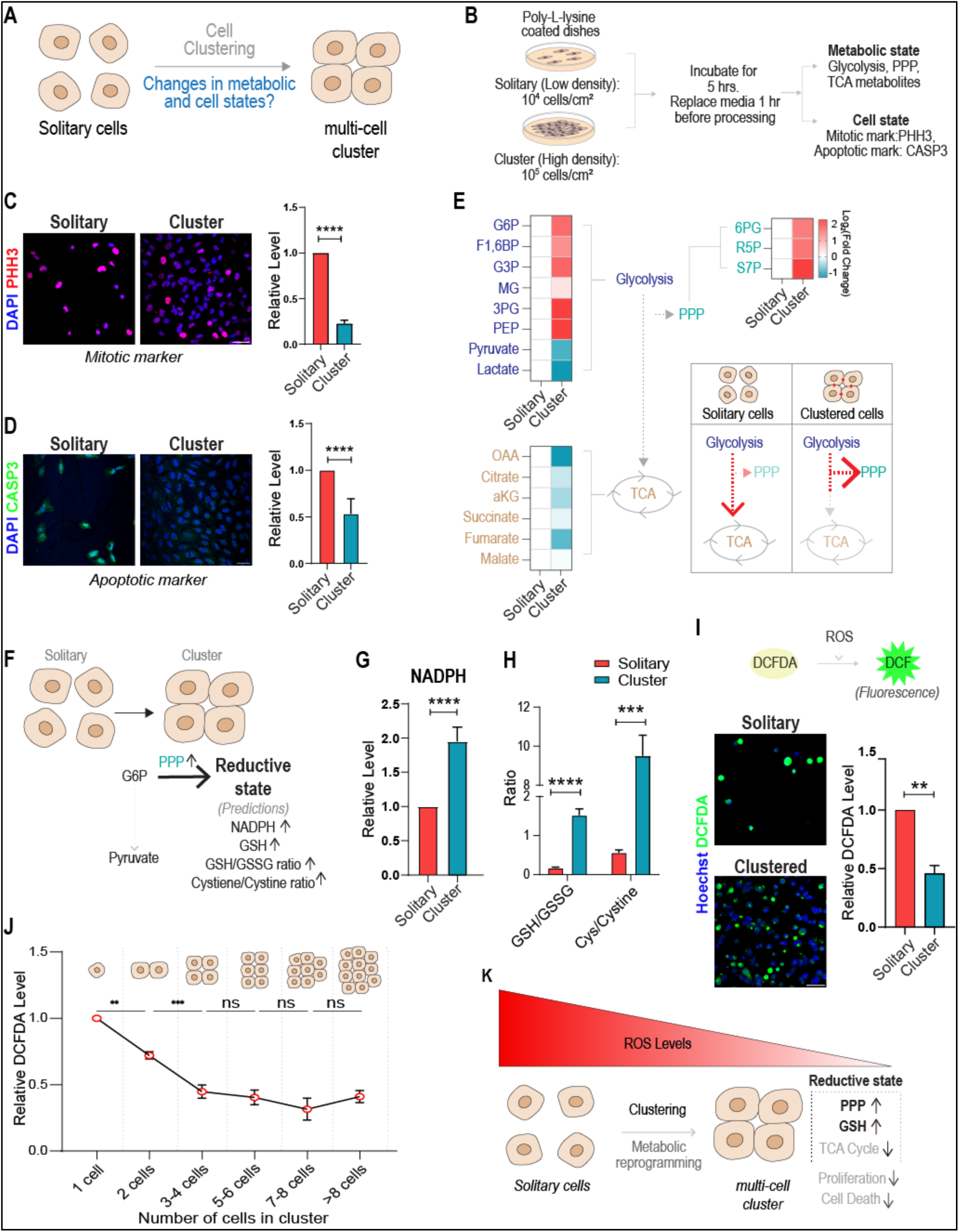
Cell clustering drives a reductive metabolic program with enhanced PPP flux and altered cell states. (A) A schematic figure, illustrating a solitary to clustered cell transition, where any early changes in the metabolic and functional state of the cell change upon cell clustering remain unknown. (B) Experimental design to address metabolic and cell state changes early as cells form clusters. Cells were seeded in low (**solitary cells**) and high density (**cell clusters**) followed by incubation for ∼5 hours, culture medium was replaced (1h), and cells were processed for quantitative metabolomics and immunocytochemistry. (C) Cell state changes in clusters of cells: Immunocytochemistry image showing the expression of phosphorylated histone 3 (PHH3), a mitotic marker, in clustered cells compared to solitary U2OS cells. Bar graph shows the relative change in the expression of PHH3. (N=3, number of cells. Solitary cells (n) = 478. Clustered cells (n) = 1611). (D) Caspase 3 marks in solitary or clustered cells: Immunocytochemistry images showing the expression of activated caspase 3 (Act. CASP3) in clustered cells compared to solitary U2OS cells. Bar graph shows the relative change in the expression of Act. CASP3. (N=3, number of cells - Solitary cells (n) = 730. Clustered cells (n) = 1458). (E) Metabolic reprogramming in cell clusters towards increased PPP: Heat map illustrates the relative changes in metabolites of glycolysis, the pentose phosphate pathway (PPP), and the tricarboxylic acid (TCA) cycle, in cells in high-density clusters, compared to solitary, low-density cells. The inset schematic summarizes the observed changes in the metabolic state of the respective cells. (N=3). (F) Metabolic reprogramming to a reductive state. The schematic synthesizes key predictions of a reprogramming towards increased PPP outputs. Cells attain a reductive metabolic state associated with increased PPP flux, which will be predicted to have increased NADPH pools that support reductive synthesis, along with increased glutathione and associated precursor amino acid synthesis, all of which increase as cells shift to a reductive state. (G) Relative NADPH levels in solitary vs clustered cells, as estimated by quantitative LC/MS/MS. (N=3) (H) Clustered cells have enhanced reduced glutathione and cysteine: Ratios reduced: oxidized glutathione (GSH:GSSG) and cysteine:cystine in solitary vs clustered U2OS cells. Solitary cell metabolite levels are normalized to 1. (N=3) (I) Clustered cells have reduced ROS levels: ROS levels in cells were detected using DCFDA staining, where cells become fluorescent due to the deacylation of DCFDA to DCF, and oxidation of DCF to fluorescent DCF by ROS. The DCFDA fluorescence in clustered cells was compared to solitary U2OS cells. Bar graph shows relative change in the levels of DCFDA (Control normalized to 1). (N=3, Solitary (n) = 984, Clustered (n) = 1888). (J) Changes in cellular ROS based on the number of cells clustering together: The line graph shows relative levels of DCFDA staining as U2OS cells go from solitary units to clusters with an increasing number of cells. Control with cluster size of 1 cell is normalized to 1. As cells go from just one cell to a cluster of two or more cells, ROS levels substantially decrease. (N=3) (L) A summary illustrating the metabolic reprogramming (and associated cell state changes) in clustered cells compared to solitary cells. Clustered cells have increased PPP, a shift to a reductive metabolic state, and associated decreases in related metabolism. Clustered cells also show reduced phospho-histone and cell death marks. \*\**p* ≤ 0.01, \*\*\**p* ≤ 0.001, \*\*\*\**p* ≤ 0.0001, ns=not significant (student’s t test). All graphs are presented as mean ± SEM.

We first characterized gross cell state changes upon cell clustering to identify the qualitative differences in the same. We therefore initially assessed the status of cell proliferation and cell death markers by immunostaining cells with phospho-histone H3 (PHH3) (22), an established marker for active mitosis, and activated caspase 3 (act. CASP3) (23), a cysteine protease that is a widely used marker for apoptosis. Interestingly, our data revealed a substantial decrease in the expression of both PHH3 (Fig. 1C) and activated CASP3 (Fig. 1D) in clustered cells, suggesting that cell clustering leads to a substantial reduction in cell proliferation and apoptotic marks. We additionally observed an increase in the expression of yH2AX, a DNA damage marker (24, 25), in solitary cells compared to clustered cells (Fig. S1C). These data collectively indicate higher marks of cell damage and death in solitary cells compared to clustered cells.

We next assessed the metabolic states of these solitary or clustered cells. To identify the changes in cellular metabolic states, we performed quantitative LC/MS/MS based metabolomic assessments from extracts of clustered or solitary U2OS cells, estimating the levels of metabolites associated with glycolysis, the tricarboxylic acid cycle (TCA), and the pentose phosphate pathway (PPP) pathways. These data revealed increased amounts of PPP metabolites, along with a smaller decrease in the TCA cycle metabolites in clustered cells compared to solitary cells (Figure 1E). This was further supported by increased amounts of metabolites of the upper glycolysis arm, and a decrease in metabolites of the lower glycolysis in clustered cells (Figure 1E). We additionally tested if the observed metabolic changes were associated with any changes in glucose uptake. For this, we cultured cells in the presence of 2-NBDG, a fluorescent glucose analogue (26), and measured staining intensities in solitary and clustered cells. Notably, we did not observe any significant changes in the intensities of 2NBDG in solitary and clustered cells (Fig. S1D).

Collectively, these data indicate that clustered cells exhibit a metabolic rewiring, with increased allocations towards the PPP. A key aspect of increased PPP flux is the enhanced generation of reducing equivalents in the form of NADPH, enabling cells to shift to what can be defined as a reductive state that augments reductive biosynthesis, and helps cells maintain redox balance (30–32) (Fig. 1F). A prediction from this metabolic reprogramming would be increased available NADPH, along with greater amounts of (reduced and total) glutathione, and precursor amino acids (Fig. 1F, Fig. S1E). We therefore next assessed and compared NADPH amounts in solitary vs clustered cells. NADPH amounts substantially increased in clustered cells, compared to solitary cells (Fig. 1G). We further assessed if clustering led to an increase in the metabolites associated with glutathione biosynthesis and a more reductive state. We observed a significant increase in the generation of glutathione and the metabolites associated with glutathione biosynthesis in clustered cells (Fig. S1F). We also measured the ratio of reduced (GSH) to oxidized (GSSG) glutathione and the cysteine to cystine ratio. An increase in GSH/GSSG and the Cysteine/Cystine ratio is an end-point indicator of a reductive state (33, 34). Notably, we also observed substantially increased GSH/GSSG and cysteine/cystine ratios in the clustered U2OS cells compared to solitary cells (Fig. 1H). Taken together, these results reveal that cell clustering is characterized by striking metabolic rewiring, alongside cell state changes (Fig. 1F). This is characterized by an increase in the generation of PPP metabolites and a shift to a reductive state, alongside a decrease in cell proliferation and cell death (Fig. 1F).

Such a reductive state in clustered cells would be expected to protect against reactive oxygen species (ROS). We therefore hypothesized that solitary cells might have higher inherent intracellular ROS, and that cell clustering would lead to a reduction in the levels of intracellular ROS because of this metabolic rewiring. To determine relative changes in the levels of intracellular ROS, we stained solitary and clustered cells with DCFDA (2’,7’-dichlorofluorescein diacetate), a fluorogenic probe commonly used for detecting ROS (35, 36). Notably, we found that solitary cells had high relative ROS levels, and cell clustering led to a significant reduction in the levels of intracellular ROS (Fig. 1I).

We next asked how dependent this reduced intracellular levels of ROS was on the formation of a cluster. To test this, we first seeded cells in increasing densities and measured DCFDA levels after 6 hours of seeding (with the experimental setup described in Fig. 1B). We observed a sharp decrease in intracellular ROS levels with increasing cell density (Fig. S1G). We therefore specifically asked if this was dependent on cells forming clusters, assessing what were the minimum number of cells that needed to come together to observe decreased ROS. To test this, we quantified DCFDA levels in cells with different cluster sizes, and compared them with ROS levels in solitary cells. Notably, we observed a significant reduction in ROS as solitary cells transitioned to just two-cell clusters (Fig. 1J). Subsequent increases in cluster size (>3 cells) led to a small further decrease in ROS, which stabilized by 5-6 cell clusters (Fig. 1J).

Summarizing, these results reveal that as solitary cells form clusters, this is characterized by a metabolic reprogramming with an increased flux through the PPP, a shift to a reductive state, and a decrease in cell proliferation and cell death (Fig. 1K). This corresponds with a sharp attenuation of intracellular ROS (Fig. 1K).

### Cadherin junction-dependent cell clustering mediates the reductive metabolic program and cell state changes

The formation of adherence junctions is one of the initial events that stabilizes cell clusters (Fig. S2A), and precedes the formation of cell-ECM junctions (Fig. S2A). We therefore next asked if the formation of adherens junctions is associated with the metabolic and cell state changes as solitary cells form clusters (Fig. 2A). Since U2OS cells form N-cadherin junctions (27), we treated the clustered cells with ADH1-trifluoroacetate (hereafter ADH1), which is an established, specific, competitive inhibitor of N-cadherin alongside DMSO vehicle controls (Fig. 2B) (21, 28, 29). Fig. 2C shows the formation of N-cadherin junctions in these cells as they form clusters. Importantly, we maintained an identical seeding density in this condition to ensure that both control and treated samples had similar cell densities and clustered cells. Treating clustered cells with ADH1 disrupted cadherin junctions, as indicated by the tapered appearance of N-cadherin immunostaining in treated cells (Fig 2D), and the clusters (with intact or disrupted junctions) are similar in scale (Fig. 2D). As additional controls, we established that treatment with ADH1 did not change cell appearance or cell size, and we did not observe any change in the nuclear area (Fig. S2B and C).

**Figure 2:**
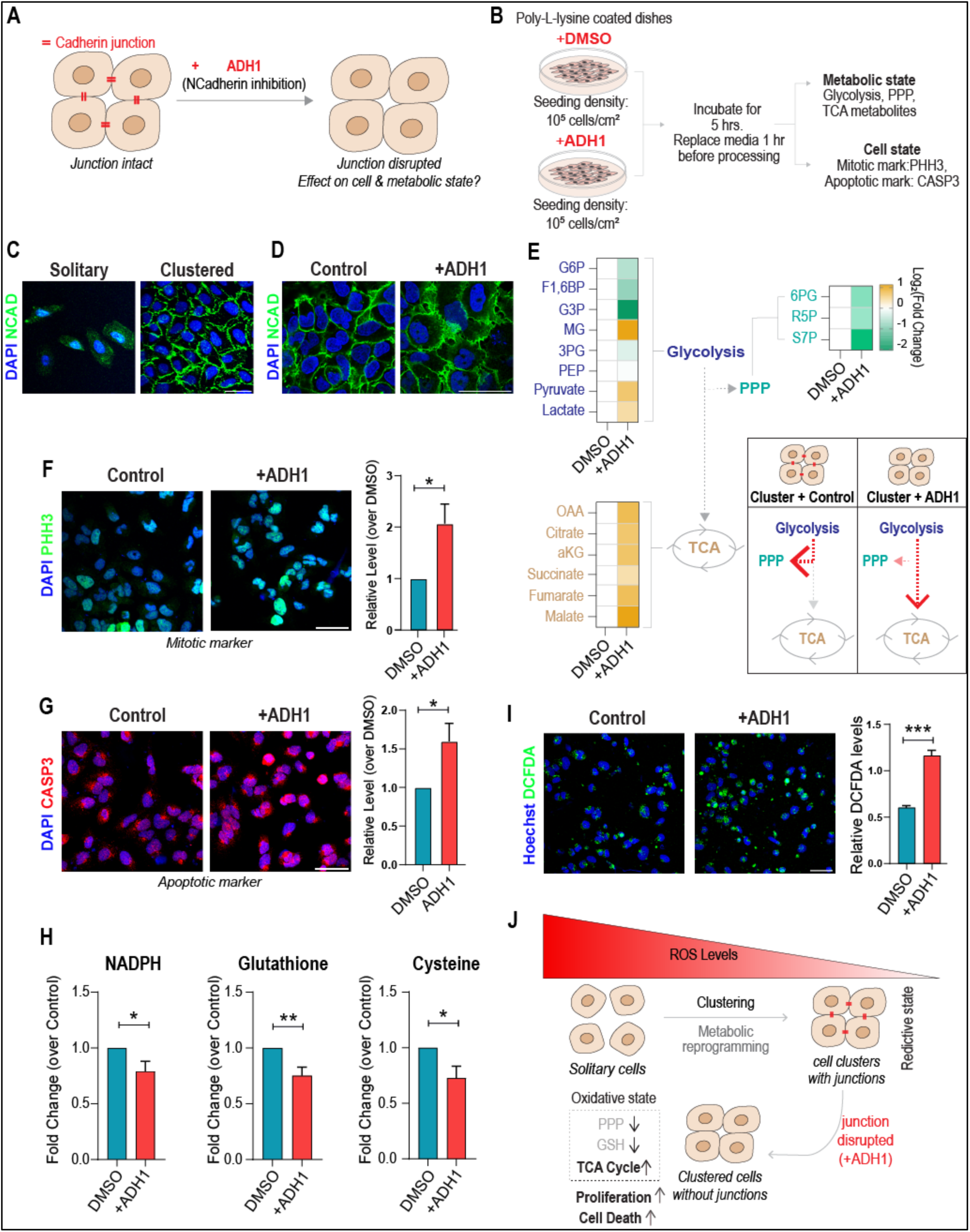
Cadherin-dependent cell clustering mediates the reductive metabolic program and cell state changes. (A) Cadherin junctions enable the formation of stable cell clusters. The cadherin junctions can be disrupted by blocking using the N-Cadherin-specific inhibitor ADH1. How does the metabolic and cell states change when cadherin junction formation is blocked in cell clusters? (B) Experimental design for LC/MS/MS-based metabolomics and cell state analysis in clustered cells, or clustered cells where Cadherin junctions are disrupted by ADH1. Cells were seeded in high-density conditions with the N-cadherin inhibitor ADH1 or DMSO control, followed by incubation for 5 hours, and replacement of cell culture medium (1 hr), and cells were processed for metabolomics or immunocytochemistry. See Methods for details. (C) N-Cadherin immunocytochemistry using an N-Cadherin-specific antibody, in solitary cells (left panel) or clustered cells (right panel), in U2OS cells (representative picture, (N=3)). (D) N-Cadherin immunocytochemistry using an N-Cadherin-specific antibody, in clustered cells (left panel), or clustered cells treated with ADH1 (right panel). The green color shows the status of the cadherin junctions in U2OS cells (N=3). (E) N-cadherin inhibition reverses the metabolic program in clustered U2OS cells to resemble solitary cells: Heat map showing the relative change in the steady-state metabolite levels of glycolysis, PPP, and TCA cycle in clustered U2OS cells after treatment with cadherin junction inhibitor ADH1 (compared to clustered cells with intact Cadherin junctions). The schematic summarizes the observed changes in the metabolic state of the clustered cells with disrupted cadherin junctions, which now show reduced PPP metabolites, and an increased glycolytic state. (N=3). (F) Cell state changes in clusters of cells with N-cadherin inhibited: Immunocytochemistry image showing the expression of phosphorylated histone 3 (PHH3) in clustered U2OS cells after ADH1 treatment compared to the control. Bar graph shows relative change in the expression of PHH3 (Control normalized to 1). (N=3, Clustered + Control (n) = 1036, Clustered + ADH1 (n) = 978). (G) Caspase 3 marks in clustered cells with N-cadherin inhibited: Immunocytochemistry image showing the expression of activated caspase 3 (Act. CASP3) in clustered U2OS cells after ADH1 treatment compared to the control. Bar graph shows relative change in the expression of Act. CASP3 (Control normalized to 1). (N=3, Clustered + Control (n) = 1581, Clustered + ADH1 (n) = 1094). (H) Reduction in levels of NADPH, glutathione (reduced), and cysteine in clustered U2OS cells upon treatment with the N-cadherin inhibitor ADH1. (N=3) (I) Effect of N-cadherin inhibition on ROS levels in clustered cells: DCFDA staining in clustered U2OS cells treated with ADH1 compared to the control. Bar graph shows the relative change in the levels of DCFDA (Control normalized to 1). (N=3, Clustered + DMSO (n) = 765 cells, Clustered + ADH1 (n) = 630 cells). (J) A summary of metabolic and cell state changes when N-cadherin is inhibited as cell clusters form. Cells with inhibited N-cadherin junctions have decreased PPP, reduced NADPH, glutathione, and increased associated carbon metabolism, cell proliferation and cell death marks, alongside increased ROS levels. Disrupting cadherin junction results in clustered cells now phenocopying low-density, solitary cells. \**p* ≤ 0.05, \*\**p* ≤ 0.01, \*\*\**p* ≤ 0.001 (student’s t test). All graphs are presented as mean ± SEM.

To assess the metabolic state of the clustered cells with perturbed junctions (ADH1 treated), we quantitatively estimated metabolites of the glycolysis, TCA, and PPP pathways as done previously. We observed that ADH1 treatment of clustered cells led to a substantial reduction in PPP intermediates, and an increase in TCA cycle metabolites (Fig. 2E). This was further supported by an increased metabolites of lower glycolysis and decreased metabolites of upper glycolysis (Fig. 2E). As a control, we did not observe any significant difference in the relative levels of ATP, suggesting that perturbation of cell-cell junctions does not change gross bioenergetics in the clustered cells (Fig. S2D). Notably, the metabolic state of clustered cells with disrupted cadherin junctions therefore closely resembled the metabolic state of the solitary cells, as defined previously in Fig. 1E. These data suggest that disrupting cell junctions in clustered cells reverts them to a metabolic state similar to solitary cells.

We next assessed the cell state changes in clustered cells treated with ADH1. To test this, we stained and quantified the expression of PHH3 and activated CASP3 in cells treated with ADH1 compared to controls, as done previously. We observed that the treatment of clustered cells with ADH1 led to a significant increase in the expression of PHH3, activated CASP3, and yH2AX marks (Fig. 2F, G and S2E). Phenotypically, the observed cell state (after disrupting cadherin junctions) again corresponds to that of solitary cells. We therefore asked if disrupting cadherin junctions in clustered cells now reversed the reductive metabolic program and its associated changes in intracellular ROS. To determine changes in the levels of intracellular ROS, we stained clustered cells with or without ADH1 treatment with DCFDA (Fig. 2I). Notably, clustered cells treated with ADH1 (with disrupted cadherin junctions) showed a striking increase in ROS levels (Fig. 2I), reversing the decrease in ROS due to cell clustering. We next addressed the state of the reductive program in clustered cells after ADH1 treatment, by assessing NADPH as well as associated glutathione and cysteine levels. Notably, disrupting the cadherin junctions using ADH1 led to a significant reduction in steady-state levels of NADPH, glutathione and cysteine (Fig. 2H). We further asked if the reduced intracellular levels of ROS as cells clustered was on cadherin junction. To test this, we seeded cells in increasing densities, with or without ADH1 treatment, and measured DCFDA levels (as described earlier in Fig. 1I, J). The sharp decrease in intracellular ROS levels observed with increasing cell density was recapitulated (Fig. S2F), however treating cells with ADH1 now attenuated this decrease. When ROS levels in clustered cells were quantified, starting with solitary cells and measured over different cluster cell numbers, the ADH1-treated cell clusters did not show a decrease in ROS (Fig. S2G), effectively reverting to a state resembling that of solitary cells.

These results reveal that the cadherin junctions are required to reprogram clustered cells towards a reductive state (Fig. 2J). The formation of cadherin junctions within a cell cluster controls the metabolic state, internal ROS levels, and cell-state changes observed in the newly clustered cells. Notably, the loss of cadherin junctions reverses the reductive program, resulting in clustered cells that resemble solitary cells in metabolic and cell states (Fig. 2J). These data collectively establish a causal role for cadherin junctions in driving cell cluster metabolic states.

### Cadherin-dependent reprogramming towards increased PPP and a reductive state is conserved in multiple cell lines

Our results in the U2OS carcinoma cell line reveal that the formation of cadherin junctions in stable cell clusters leads to a reductive metabolic reprogramming that restrains intracellular ROS levels. We asked if this cadherin-dependent metabolic reprogramming, and the associated ROS reduction, is recapitulated in multiple other, non-cancerous cell lines. To test this, we assessed distinct cells types - mouse embryonic fibroblasts (MEF SV40 and NIH3T3) and Human foreskin fibroblasts (BJhTERT), and measured the relative levels of ROS using DCFDA. Notably, cell clustering led to a ubiquitous reduction in cellular ROS levels as suggested by reduced intensity of DCFDA stain in these cell lines (Fig. S3A and B). We next asked if the clustering-dependent reduction in ROS is associated with an increased PPP pathway flux and the corresponding reductive metabolic program. To test this using mouse embryonic fibroblasts (MEF), we measured steady-state levels of the metabolic pathway (as assessed previously) in solitary and clustered MEFs. We observed that cell clustering in MEFs leads to a striking increase in upper glycolysis and PPP metabolites (Fig. S3C and D). Furthermore, we also observed increased levels of NADPH and metabolites associated with glutathione synthesis and reductive metabolism (Fig. S3E and F). Finally, we assessed the dependence of this reductive program on the cadherin junction. Treatment of clustered MEFs with ADH1 (the cadherin junction inhibitor) led to a decrease in upper glycolysis, PPP, NADPH, glutathione, and cystine metabolism (Fig. S3G-I). Together, these data indicate a conserved role for adherence junctions in driving PPP and reductive metabolism in non-cancerous cell lines. Collectively, these results suggest that mammalian cells cluster and form cadherin junctions, and ubiquitously shift to a reductive metabolic program with enhanced PPP flux, and restrain internal ROS.

### Cadherin junction dependent metabolic reprogramming enhance the resistance of clustered cells to exogenous ROS

Clustered cells form cadherin junctions, which reprogram their metabolism towards a reductive state, resulting in reduced intracellular ROS levels. We therefore asked if solitary and clustered cells differ in their ability to resist the effect of exogenous ROS inducers, as illustrated in Fig. 3A. To increase ROS pharmacologically, we treated the cells with Erastin, a ROS inducer that depletes intracellular cystine by decreasing uptake and, in turn, reduces glutathione to increase ROS (37). Notably, erastin treatment led to a further increase in ROS in solitary cells, but did not do so in clustered cells (Fig. 3B), indicating that clustered cells show increased resistance to external ROS inducers. This increased ROS in solitary cells led to a further increase in the PHH3 and activated caspase 3 markers in solitary cells (Fig. 3C and D). Notably, in clustered cells the erastin treatment did not lead to further changes in these associated cell state markers (Fig. 3C and D). Together, these results indicate that clustered cells are buffered against external ROS stressors, further reinforcing the outcome of their shift to a reductive state.

**Figure 3:**
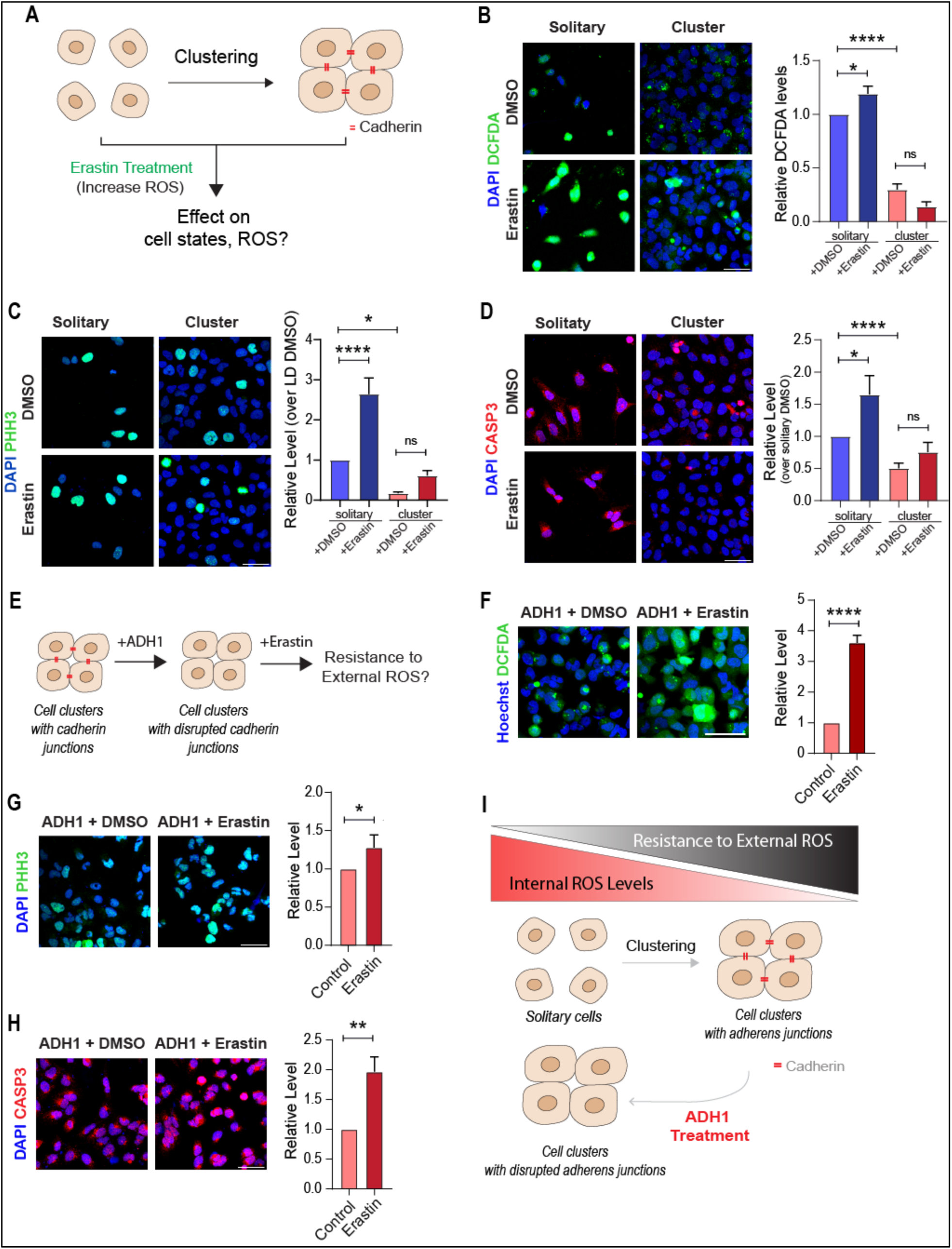
Cadherin junction dependent metabolic reprogramming enhances the resistance of clustered cells to exogenous ROS. (A) Experimental design to assess changes in intracellular ROS and cell states of solitary and clustered cells after treatment with a strong ROS inducer - Erastin. (B) Effect of Erastin treatment on intracellular ROS in solitary or clustered U2OS cells: Images showing DCFDA (ROS indicator) levels in solitary and clustered cells treated with Erastin and DMSO control. Bar graph shows the relative change in the expression of DCFDA. Staining intensity of solitary cells normalized to 1. (N=3, Solitary + DMSO (n) = 809, Solitary + Era (n) = 706, clustered + DMSO (n) = 3179, clustered + Era (n) = 3123). (C) Cell state changes in Erastin-treated solitary and clustered U2OS cells: Immunocytochemistry images showing relative expression of PHH3 in solitary and clustered cells treated with Erastin and DMSO control. Bar graph shows the relative change in the expression of PHH3. Staining intensity of solitary cells normalized to 1. (N=3, Solitary + DMSO (n) = 690, Solitary + Era (n) = 703, clustered + DMSO (n) = 3772, clustered + Era (n) = 3950). (D) Caspase 3 marks after Erastin treatment in solitary and clustered U2OS cells: Immunocytochemistry images showing relative expression of activated CASP3 in solitary and clustered cells treated with Erastin and DMSO control. Bar graph shows the relative change in the expression of activated CASP3. Staining intensity of solitary cells normalized to 1. (N=3, Solitary + DMSO (n) = 613, Solitary + Era (n) = 629, clustered + DMSO (n) = 3758, clustered + Era (n) = 2167). (E) Experimental design to assess the response of clustered cells to ROS inducer Erastin after cadherin disruption (by adding ADH1). (F) ROS levels in clustered cells with N-cadherin inhibition and ROS induction: Images showing DCFDA (ROS indicator) levels in ADH1-treated cells further treated with Erastin and DMSO control. Bar graph shows the relative change in the levels of DCFDA staining. (N=3, ADH1 + DMSO (n) = 4127, ADH1+Era (n) = 3814). (G) Cell state changes in Erastin-treated clustered cells with N-cadherin inhibition: Immunocytochemistry images showing relative expression of PHH3 in ADH1-treated cells further treated with Erastin and DMSO control (N=3, ADH1 + DMSO (n) = 3645, ADH1+Era (n) = 4360). (H) Caspase 3 changes in Erastin-treated clustered cells with N-cadherin inhibition: Immunocytochemistry images showing relative expression of activated CASP3 in ADH1-treated cells further treated with Erastin and DMSO control (N=3, ADH1 + DMSO (n) = 4468, ADH1+Era (n) = 4674). (I) Schematic illustrating how clustered cells form cadherin junctions to become resistant to the effects of ROS induction. The disruption of cadherin junctions results in cells becoming sensitive to ROS induction and phenocopy solitary cells. \**p* ≤ 0.05, \*\**p* ≤ 0.01, \*\*\*\**p* ≤ 0.0001, ns=not significant (student’s t test). All graphs are presented as mean ± SEM.

We next asked if this increased resistance to exogenous ROS was cadherin-dependent, by disrupting cadherin junctions and assessing the sensitivity of clustered cells to ROS induced by erastin (Fig. 3E). To test this, we treated clustered cells with junction inhibitor ADH1 (as described earlier) and added the ROS inducer erastin. Notably, ADH1 treatment led to an increase in ROS levels in clustered cells with ADH1 treatment after erastin treatment (Fig. 3F). Clustered ADH1-treated cells when subjected to erastin also showed further increase in cell proliferation and cell death markers (Fig. 3G and H). Effectively, the ADH1-treated clustered cells subject to ROS generated by erastin now phenocopied the cellular response shown of solitary cells.

Taken together, these results reveal that clustered cells stabilized by the cadherin junctions have enhanced resistance to external ROS inducers (Fig. 3I), due to their cadherin-dependent reprogramming to a reductive state.

### Inhibiting PPP flux sensitizes clustered cells to ROS and exacerbates ROS-induced cell state changes

Since cadherin-mediated cell clustering leads to increased PPP flux and NADPH, and restrains intracellular ROS levels, we next addressed the consequences of inhibiting PPP in clustered cells (with intact junctions) on intracellular ROS and cell states (Fig. 4A). To inhibit PPP, we utilized specific inhibitors of PPP flux, namely G6PDi-1 and Polydatin. These inhibitors block glucose-6-phosphate-dehydrogenase (G6PDH), the enzyme catalyzing the first, critical step of the oxidative PPP arm, as shown in the schematic (38–40) (Fig. 4B). Inhibiting PPP using G6PDi-1 or polydatin led to an increase in ROS in clustered cells with cadherin junctions (Fig. 4C). This reiterates the requirement of the metabolic programming towards increased PPP flux to retain a reductive state. We next assessed the cell state markers after inhibiting the PPP in clustered cells with cadherin junctions. PPP inhibition led to decreased cell proliferation marks in these cells (Fig. 4D), which is potentially due to a critical role for the PPP pathway in the synthesis of nucleic acids necessary for cell proliferation. PPP inhibition also led to an increase in the expression of activated CASP3 (Fig. 4E). We further asked if PPP inhibition led to an increase in ROS in clustered cells treated with ADH1. The treatment with PPP inhibitors and ADH1 led to an increase in intracellular ROS and cell death (Fig. S4A-C), reiterating the central role for PPP metabolism in maintaining cellular redox states. We next asked if PPP inhibition alone exacerbates the sensitivity of clustered cells to exogenous ROS and erastin-induced cell death, as observed with solitary cells or cadherin disruption in cell clusters (Fig. 4F). To test this, we treated clustered cells with PPP inhibitor G6PDi-1 and erastin. The controls were treated with DMSO. As expected, erastin treatment led to an increase in intracellular ROS, a reduction in proliferating markers, and an increase in cell death markers in PPP-inhibited clustered cells compared to the controls (Fig. 4G, 4H).

**Figure 4:**
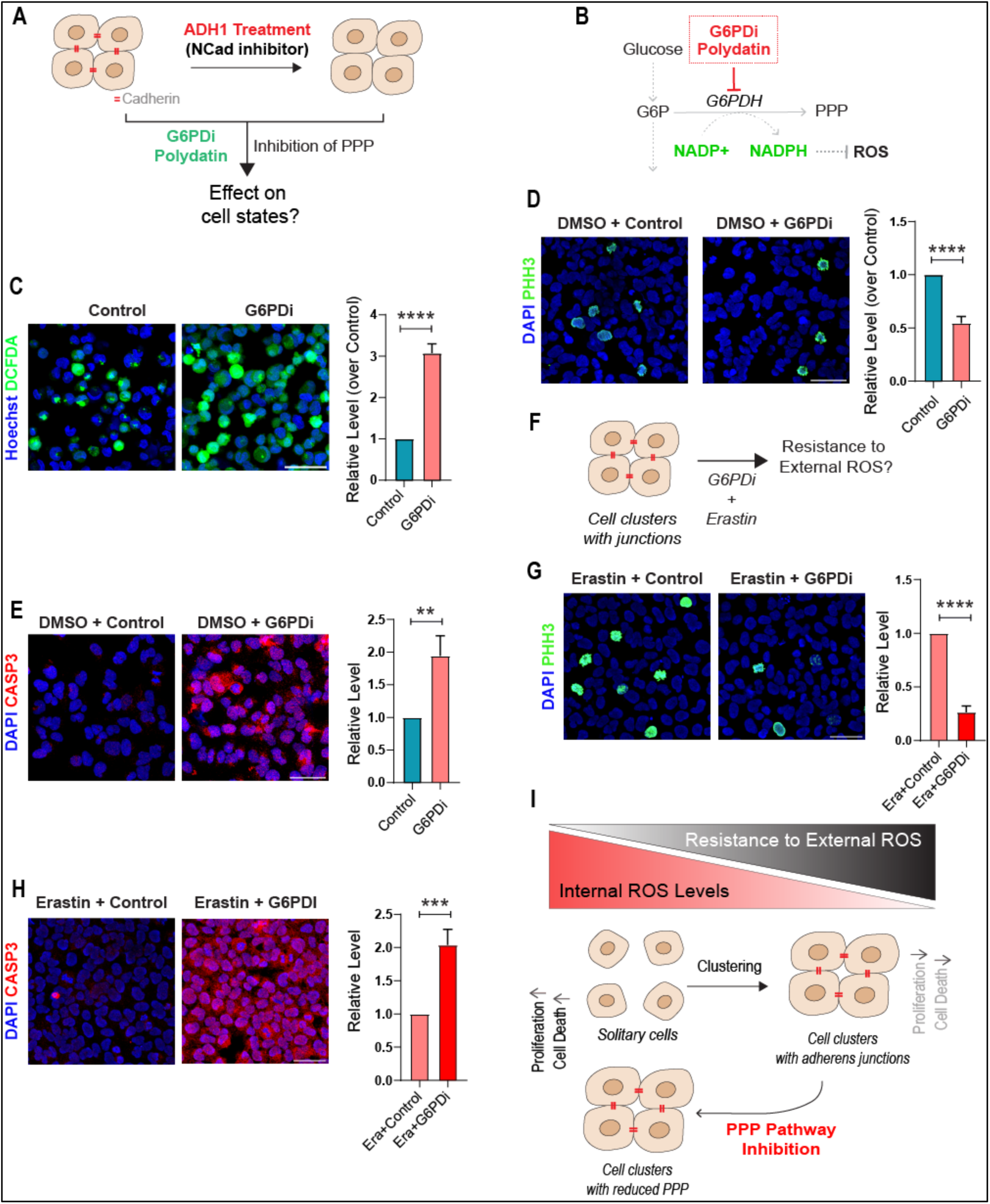
Inhibiting PPP flux sensitizes clustered cells to ROS and exacerbates ROS-induced cell state changes. (A) Experimental design to assess the cell states of clustered cells, with and without cadherin junctions, after treatment with inhibitors of the PPP pathway. (B) Schematic showing the specific inhibitors of the PPP used (G6PDi-1, Polydatin and 2DG), also indicating the nodes in the PPP that are inhibited by G6PDi-1, Polydatin and 2DG. (C) ROS levels in clustered cells with the PPP inhibited: Image showing expression of DCFDA in clustered cells after treatment with G6PDi-1 and DMSO control (N=3, Cluster + Control (n) = 2938, Cluster + G6PDi (n) = 3358). (D) Cell state changes in clustered cells upon inhibiting the PPP: Immunocytochemistry image showing the expression of PHH3 in clustered cells treated with G6PDi-1 (N=3, Cluster + Control (n) = 1022, Cluster + G6PDi (n) = 1076). (E) Caspase 3 marks in clustered cells upon inhibiting the PPP: Immunocytochemistry image showing the expression of activated CASP3 in clustered cells treated with G6PDi-1 (N=3, Cluster + Control (n) = 1248, Cluster + G6PDi (n) = 1477). (F) Experimental design to assess response to ROS and changes in cell states in clustered cells when treated with PPP inhibitor G6PDi-1, and ROS inducer Erastin. (G) Cell state changes in clustered cells after Erastin and PPP inhibitor treatment: Immunocytochemistry image showing expression of PHH3 in clustered cells after treatment with PPP inhibitor G6PDi-1 and ROS inducer Erastin, and control (N=3, Era + Control (n) = 1415, Era + G6PDi (n) = 1741). (H) Caspase 3 marks in clustered cells after Erastin and PPP inhibitor treatment: Immunocytochemistry image showing expression of activated CASP3 in clustered cells after treatment with PPP inhibitor G6PDi-1 and ROS inducer Erastin and control (N=3, Era + Control (n) = 1436, Era + G6PDi (n) = 2117). (I) Schematic illustrating how clustered cells with cadherin junctions maintain reduced ROS and a reductive state via increased PPP. Disrupting cadherin junctions, or inhibiting only the PPP flux, results in high ROS and associated cell state changes. \*\**p* ≤ 0.01, \*\*\**p* ≤ 0.001, \*\*\*\**p* ≤ 0.0001 (student’s t test). All graphs are presented as mean ± SEM.

Taken together, these results reveal that the augmented PPP flux in clustered cells with stable cadherin junctions enables cells to maintain a reductive state, restrain internal ROS, and resist the effects of exogenous ROS, while concurrently controlling cell states (Fig. 4I).

### Cadherins mediate the reductive program in cell clusters by activating NRF2

Finally, we addressed the molecular mechanisms that drive the reductive metabolic program due to the cadherin-mediated junctions formed in clustered cells (Figure 5A). The NRF2 signaling pathway (mediated by the NRF2 transcriptional regulator) is a primary regulator of reductive programs in mammalian cells, including an increase in PPP metabolism, NADPH, and glutathione metabolism (41). We therefore assessed the status of the NRF2 pathway in cell clusters and its dependence on cadherin junctions. To test this, we first determined the relative expression of NRF2 protein in control and ADH1-treated cells at high cell density (Fig. 5B). Inhibiting cell junction formation in the clustered cells using ADH1 led to a rapid decrease in NRF2 protein expression, observed as early as 3 hours of inhibitor treatment, establishing that this is an early response to junction formation (Fig. 5B). We next assessed the expression of transcripts of NFR2, KEAP1 and other downstream targets of the NRF2 pathway, and observed a substantial decrease in these transcripts in ADH1-treated cells (Figure 5C). Together, these results indicate that cadherin junction formation in clustered cells increases NRF2 and NRF2-dependent outputs.

**Figure 5:**
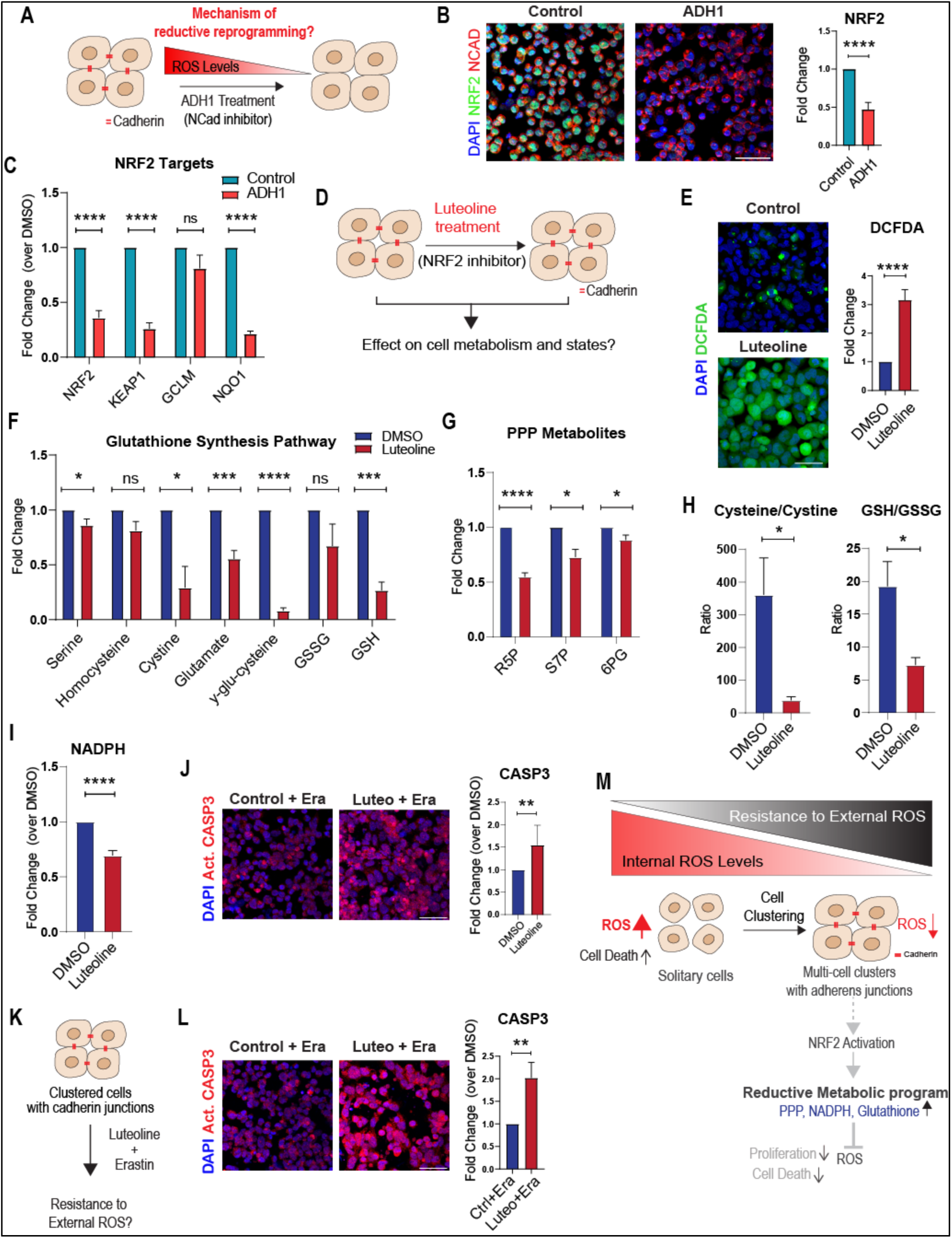
Cadherins mediate the reductive program in cell clusters by activating NRF2. (A) Cell clustering and the formation of the cadherin junction leads to a reductive metabolic reprogramming that restrains intracellular ROS levels in clustered cells. This experimental setup allows us to ask about the mechanism by which cadherins control this reductive metabolic reprogramming. (B) NRF2 expression in clustered cells is cadherin dependent: The figure shows NRF2 expression in clustered cells before or after ADH1 treatment: immunocytochemistry image showing expression of NRF2 protein after 3 hours of treatment with ADH1. The bar graph on the right shows the relative expression of NRF2 proteins in ADH1-treated clustered cells compared to controls (N=3, Cluster + DMSO (n) = 2573, Cluster + ADH1 (n) = 2407). (C) Graph showing relative levels of NRF2 pathway-dependent transcripts in ADH1-treated cells compared to the controls after 6 hours of treatment with ADH1, as measured using RT-qPCR (N=3). (D) Schematic illustrating the experimental design to test how the inhibition of NRF2 using Luteoline impacts metabolism and cell states in clustered cells. (E) DCFDA staining showing the levels of intracellular ROS in Luteoline-treated clustered cells compared to the controls. The bar graph on the right shows the relative levels of DCFDA in luteoline-treated clustered cells compared to controls (N=3, Cluster + DMSO (n) = 5221, Cluster + Luteoline (n) = 1141). (F) Metabolomics data showing relative levels of metabolites associated with glutathione metabolism in the clustered cells treated with luteoline compared to controls (N=3). (G) Metabolomics data showing relative levels of metabolites associated with PPP metabolism in the clustered cells treated with luteoline compared to controls (N=3). (H) Metabolomics data showing ratios of GSH/GSSG and cysteine/cystine in clustered cells treated with luteoline compared to controls. The data reflects on the redox state of the clustered cells after luteoline treatment. (N=3). (I) Relative NADPH levels in clustered cells treated with luteoline compared to controls (N=3). (J) CASP3 marks on clustered cells without or with Luteoline treatment: Immunocytochemistry image showing expression of activated CASP3 in clustered cells after treatment with NRF2 inhibitor, Luteoline inhibitor, and control. The bar graph on the right shows relative levels of activated CASP3 in Luteoline-treated clustered cells compared to the control (N=3, Cluster + DMSO (n) = 2910, Cluster + Luteoline (n) = 3241). (K) Experimental design to address how clustered cells wherein NRF2 is inhibited using Luteoline respond to extracellular ROS induced by Erastin. (L) CASP3 marks on clustered cells treated with Luteoline and Erastin: Immunocytochemistry image showing expression of activated CASP3 in clustered cells after treatment with NRF2 inhibitor, Luteoline, and ROS inducer, Erastin, and control. The bar graph on the right shows relative levels of activated CASP3 in Luteoline and Erastin-treated clustered cells compared to the control. (N=3, Era + DMSO (n) = 1103, Era + Luteoline (n) = 1307). (L) Model illustrating the formation of multi-cell clusters mediated by Cadherin junctions, and the corresponding metabolic reprogramming. Cadherin junction formation reprograms clustered cells towards a reductive state, with high PPP flux, as mediated by the activation of NRF2. This metabolic reprogramming enables clustered cells to have lower inherent ROS, become resistant to the effects of ROS, and have low cell death. \**p* ≤ 0.05, \*\**p* ≤ 0.01, \*\*\**p* ≤ 0.001, \*\*\*\**p* ≤ 0.0001, ns=not significant (student’s t test). All graphs are presented as mean ± SEM.

We next asked if the activation of the NRF2 pathway in the clustered cells controlled the metabolic reprogramming observed in these cells (Figure 5D). To test this, we assessed the consequences of treating clustered cells with the NRF2 pathway-specific inhibitor, luteoline (Fig. 5D). Treating clustered cells with luteoline expectedly led to a decrease in the expression of NRF2 (Fig. S5A). With this experimental setup, we assessed changes in ROS and the reductive state in clustered cells. Treatment with luteoline led to a substantial increase in intracellular ROS levels as indicated by DCFDA staining (Figure 5E). This increase in ROS due to luteoline was observed in clustered cells that were treated with ADH1, reiterating a critical role for NRF2 in maintaining cellular redox states, downstream of cadherin formation (Figure S5B). We therefore asked if the increase in the intracellular ROS observed after NRF2 treatment was due to abrogating the PPP metabolic reprogramming in the clustered cells. To test this, we determined the intracellular levels of PPP intermediates, NADPH, and metabolites associated with glutathione metabolism after inhibiting NRF2. Luteoline treatment led to decreased levels of metabolites associated with PPP and glutathione metabolism (Fig. 5F and 5G). We also observed decreased NADPH, cystine, glutathione (GSH), and GSH/GSSG, and a lower cysteine/cystine ratio, suggesting a significant reduction in the reductive capacity of the clustered cells (Figure 5H, Fig. 5I), alongside associated changes in glycolysis and the TCA cycle metabolites (Figure S5C and D). Notably, these changes mirrored the earlier observed changes in the clustered cells, wherein cadherin junctions were disrupted. Together, these data establish that the formation of cadherin junctions in clustered cells activates the NRF2 pathway and reprograms them towards a reductive program via this activation.

Since ROS levels increase in luteoline-treated cells, we next asked if this also increases cell death markers in the cell clusters. Notably, inhibiting NRF2 by luteoline treatment enhances cell death marks in the clustered cells, as observed by an increase in the expression of activated caspase 3 (Fig. 5J). Finally, we asked if the treatment with luteoline increased the sensitivity of clustered cells to erastin-induced ROS (Fig. 5K). Notably, we observed a further increase in activated CASP3 expression in clustered cells treated with luteoline and erastin (Fig. 5L). These data establish that the inhibition of the NRF2 pathway in the clustered cells increases their sensitivity to external ROS inducers.

Collectively, these data reveal that cells form cluster through cadherin junctions, leading to the activation of NRF2 (Fig. 5M). This enables a metabolic reprogramming in clustered cells that augments a reductive state, characterized by enhanced PPP flux, the generation of reductive equivalents through NADPH, and increased glutathione and associated metabolites. This enables clustered cells to maintain lower intracellular ROS, enhances their resistance to exogenous ROS inducers, and decreases cell death and other cell state marks (Fig. 5M).

## Discussion

In this study, we discover that as solitary cells assemble into clusters, the cadherin junction mediates a metabolic reprogramming in these clusters. The cadherin junctions augment a reductive metabolic state, characterized by increased PPP flux, the generation of reducing equivalents (NADPH), and higher levels of reduced glutathione and associated metabolites (Fig. 5M, model). Consequently, clustered cells with cadherin junctions also restrain intracellular ROS, and can resist exogenous ROS stress. This cadherin-mediated reductive metabolic rewiring and decrease in ROS in the clustered cells also lead to reduced cell death. Notably, disrupting cadherin junctions, or inhibiting the reductive program using small molecule inhibitors of the PPP, abrogates the resistance to ROS in cell clusters and results in increased cell death marks and hypersensitivity to external ROS inducers. Mechanistically, the cadherin junctions activate NRF2 signaling, which stabilizes the reductive program. Our study thus discovers a crucial role for cadherin junction formation in metabolically reprogramming cells towards a reductive state, as cells transition from solitary individuals to clusters (Fig. 5M).

Understanding the relationship between metabolic states and ROS is important in the context of cell state control. Cells must dynamically calibrate their reductive capacity to sustain biosynthesis, maintain metabolic efficiency, and also override the deleterious outcomes of excessive ROS. Excessive generation of ROS leads to oxidation of cellular proteins and lipids and subsequent cell death, while very low ROS levels push cells into a state of dormancy. As a result, cells have regulatory mechanisms in place to maintain intracellular ROS levels in a ‘Goldilocks’ zone, retaining the signaling and metabolic functions of ROS, while maintaining their ability to carry out biosynthesis. An effective strategy employed by cells to restrain ROS is to augment PPP flux. Enhanced PPP flux generates NADPH, which provides critical reducing equivalents in the cytoplasmic environment (31). This enables cells to carry out reductive biosynthesis as required (31, 32, 42), is coupled with glutathione (and other reductants) production, and the necessary recycling of oxidized glutathione to reduced forms, to restrain excessive ROS. The increased PPP flux simultaneously generates precursors to generate nucleotides and lipids, which are coupled to NADPH utilization, and essential raw materials to sustain cellular requirements (31, 32, 42). We have uncovered here that as cells transition to stable clusters through the formation of cadherin junctions, the junctions drive this reprogramming towards an efficient, reductive state. This metabolic reprogramming in turn restrains levels of ROS, controls the cell state, and prevents its adverse effects.

The origins of multicellularity are complex, having independently evolved many times (1, 4), and adherens junction formation is a key aspect of metazoan multicellularity. This transition is thought to have facilitated co-operative interactions that increased cell survival, specialization of function, and adaptability in the metazoans (1, 4). A crucially overlooked biochemical requirement of this evolutionary transition is the primary requirement for cell groups to maintain a stable, reductive internal environment for efficient metabolism. How the transition to multicellularity affected the ability of metazoans to integrate external oxygen levels and buffer internal redox state is poorly understood. Several reports have hypothesized mechanisms that improved ROS buffering, including ROS compartmentalization in organelles, enhancement of the antioxidant defense system, and metabolic rewiring. Interestingly, the transition to metazoan multicellularity coincides with the expression of cadherin-like molecules, termed proto-cadherins, that facilitate the formation of stable cell clusters (43, 44). The formation of stable cadherin junctions may potentially be a general mechanism that allows clustered cells to shift to an efficient, reductive state, augment reductive biosynthesis, and suitably restrain intracellular ROS levels. Our results, therefore, add exciting new directions of study using experimental models of multicellularity.

It is interesting to contextualize the formation of adherens junctions with the more well-studied, ECM-mediated focal junctions. Adherence (cell-cell junctions) and focal (cell-ECM) adhesion proteins have long been studied in the context of cell fates, cell migration, tissue morphogenesis, mechanical sensing, and communication (45). Evidence in the literature also suggests a role of integrin junctions in regulating cellular metabolism (46). The integrin junctions are engaged upon attachment to the ECM, and this leads to context-dependent changes in AMP-regulated protein kinases (AMPK), mammalian target of rapamycin (mTOR), and hypoxia-induced factor (HIF-1α) (47–49), in turn affecting cellular outcomes. In contrast, the role of the primordial cell-cell junctions - which logically precede the formation of ECM-mediated junctions as cells form clusters - in setting up metabolic programs is unexplored. Our data reveal that the formation of the adherens junction reprograms cells to a reductive state, highlighting a causal role of cadherin junctions in programming metabolism and, in turn, controlling cell states. These results establish a foundation to elucidate hierarchies of metabolic outcomes due to cadherin engagement in stabilizing cell clusters. This can help unravel the temporal sequence and contribution of cadherin junctions in directing metabolic outcomes as solitary cells form cellular collectives.

We speculate that our findings, made using reductionist, first principles approaches, might conceptually inform studies in the field of metastatic cancer biology. In invasive cancers such as melanoma and ductal breast carcinoma, malignant cells are known to migrate in cell clusters (50). These malignant cancers are resistant to anti-cancer treatment, which works by increasing intracellular ROS (50–52). Recent studies on clustered cancer cells suggest that detached cancer cells regulate intracellular ROS levels by efficiently clearing damaged mitochondria, thereby reducing the intracellular ROS burden (9, 10). Our results offer a plausible explanation for this increased resistance of cancer cell clusters to ROS induction through the maintenance of strong cadherin junctions. A cadherin-dependent augmentation of a reductive metabolic program could influence situations such as anoikis due to ECM detachment, and reduce the sensitivity of cancer cells to exogenous ROS inducers, and can be tested in suitable models.

In conclusion, we uncover a fundamental function for the formation of adherens junctions (through cadherins) in metabolic state control, as cells transition from solitary individuals to stable clusters of cells. As solitary cells come together and form stable junctions through adherens junctions, the cadherins initiate a metabolic reprogramming towards a reductive state. This, in turn, controls cell states by augmenting this reductive capacity and restraining ROS. These highlight a previously unknown function of cadherin proteins in driving cell state transitions via metabolic control. Based on these results, we propose that the ability to form stable junctions played a key role in enabling cell groups to program their metabolism and maintain a reduced cellular environment. This adds an exciting dimension towards addressing the origins of multicellularity as cells formed clusters, based on a primordial need to control metabolic states in cell groups.

## Materials and Methods

### Cell culture

U2OS cells, BJhTERT, NIH3T3, and SV40 MEFs were grown in DMEM media (Sigma) with 1% Glutamax (Sigma) at 37°C for ∼6 hours. One hour prior to cell processing, the cell culture media was replaced. To maintain cells in a solitary state, cells were seeded at a low density, 10^4 cells/cm2. To promote cells clustering, cells were seeded at a high cell density 10^5 cells/cm2. The passage number of the cells varies from P11 to P18. For all the experiments, the inhibitors were added at the time of cell seeding. Vehicle (DMSO) was added in the control experiments. The list of drugs, their sources, and the concentrations used are given in Supplementary Table 1 below.

#### 1. Cell culture for immunocytochemistry and DCFDA staining

For immunocytochemistry, cells were seeded on glass plates attached to punched cell culture dishes. Glass plates were treated with poly-l-lysine (PLL). For PLL coating, the cover glass attached to the punched dishes was incubated with PLL at room temperature for 15 minutes. PLL was subsequently removed, and the excess was washed with sterile double-distilled water. Following PLL coating, cells were plated at above mentioned cell density. 1 hour prior to cell processing, the media was replaced. Cells were processed for immunocytochemistry after ∼6h.

#### 2. Cell culture for metabolomics, biochemical assays, and protein isolation

For metabolomics, protein isolation, and biochemical assays, cells were seeded in low- and high-density conditions on non-TC-treated culture plates treated with PLL. The PLL coating was done for 20 minutes, followed by washing with sterile distilled water. 1 hour prior to cell processing, the media was replaced. Cells were processed for metabolite and protein extraction after ∼6h.

### ATP assay

Cells were lysed using 1X lysis buffer (10 mM Tris (pH 7.5), 100 mM NaCl, 1 mM EDTA, 0.01% Triton X-100) for 30 - 60 mins with constant agitation at 4^ο^C, followed by centrifugation at 14000 g, 4^ο^C, 15 mins. The supernatant containing the protein fraction was collected. Protein estimation was done by BCA (bicinchoninic acid) protein assay kit (G-BioSciences, catalogue number-786-570). ATP levels were measured using the ATP estimation kit (Thermo Fisher A22066). All measurements were normalised to protein amounts detected by the BCA assay.

### Immunocytochemistry

Cells seeded on the PLL-coated glass plates attached to cell culture dishes were washed twice with 1XPBS. Fixation was done with 4% paraformaldehyde for 8 minutes and washed three times with 1XPBS. Permeabilization was done using 1XPBS with 0.5% Triton X-100 for 10 minutes. This is followed by blocking with 0.5% PBS-T with 5% normal donkey serum for 45 minutes. Primary antibody staining was done overnight. Secondary antibody staining was done for 45 minutes. Confocal imaging was done at the Central Imaging and Flow Cytometry Facility (CIFF) in the InStem facility using FV3000 5L and a resonant scanner. A list of antibodies, their sources, and concentrations is provided in Supplementary Table 2 below.

### Image processing and analysis

All images presented in the manuscript are z-stack confocal images taken in the confocal microscopes mentioned above. Laser powers and voltages were kept strictly identical for exciting the fluorophore attached to the protein of interest. Non-linear adjustments might be made for DAPI staining for improved visualization of the cells. For presentation in the manuscript, only identical linear adjustments in the image brightness and contrast have been made using ImageJ software for better visualization of the protein of interest. Non-linear adjustments might be made for the blue channel with DAPI staining for better visualization of the nucleus.

For intensity quantification, mean fluorescence intensities of all the cells in a large field of view were collected. For image analysis, a custom script was written on the ImageJ software. Final mean fluorescence intensities were calculated by taking an average of the mean fluorescence intensities of all the cells in a field of view. On average, 300-1500 cells were taken per sample to determine the average mean fluorescence intensities.

### DCFDA Staining and Quantification

2’,7’-dichlorodihydrofluorescein diacetate (DCFDA) (Cat. No. D399, Thermo Fischer Scientific) was used to detect ROS in all the cells mentioned in the manuscript. Cells cultured on the coverslip, as described above, were incubated with 10 uM concentration of DCFDA prepared in DMEM media for 30-45 minutes at 37 degrees Celsius. Excess DCFDA was washed using 1X PBS. Nucleus was stained using Hoechst-33342 (62249, Thermo Fischer Scientific) for 3 minutes. Stained cells in DMEM were imaged in the green channel in FV3000 5L.

### 2-NBDG staining for Glucose estimation

2-NBDG (2-[N-(7-nitrobenz-2-oxa-1,3-diazol-4-yl)amino]-2-deoxy-D-glucose) (N13195, Thermo Fisher Scientific), a fluorescent glucose analogue, was used as a proxy to determine the extent of glucose uptake in different conditions as described in the manuscript. 6h after seeding, cells were starved in glucose-deficient DMEM media (11966025, Thermo Fischer Scientific) for 1 hour. Cells were subsequently incubated with 2NBDG-containing DMEM media at a working concentration of 200 uM for 1 hour at 37 degrees Celsius. Cells were washed with PBS. Stained cells in the DMEM media were imaged in the green channel in FV3000 5L immediately.

### Steady state metabolomics

#### 1. Steady-state metabolomics sample preparation

Adherent cells grown on PLL-coated non-TC-treated dishes were washed with ice-cold 1X PBS twice on ice. 1 mL of cold extraction buffer (Ethanol: H_2_O, 4:1 v/v, mass spectrometry grade) was added to all dishes. Cells were quickly scraped and kept on ice. To ascertain the extent of efficiency of extraction, an internal control of 13C4–aspartate was used. Metabolite extraction was done by placing the Eppendorf tubes on ice for 1 minute, followed by vortexing for 1 minute. Samples were centrifuged at 16000g for 15 minutes to pellet cellular debris. The supernatant was divided into two tubes, one each for amino acids/sugar phosphates and unstable glycolysis/TCA cycle metabolites, directly or after derivatization using *O*-benzylhydroxylamine (OBHA), using methods developed earlier (53). The samples were subsequently vacuum dried using a SpeedVac vacuum concentrator. Samples were stored at –80°C until LC–MS/MS analysis. Prior to injection into the mass spectrometer, metabolites were resuspended in mass spectrometry-grade water. The peak intensity data were normalized to cell seeding density, and protein levels quantified by BCA.

#### 2. Metabolite profiling

Glycolysis, TCA, and amino acid metabolites were detected using quantitative LC-MS/MS as described previously (53). LC–MS/MS analyses were performed using an AB SCIEX QTRAP 5500 system connected to a Shimadzu Nexera UPLC. A Synergi 4 μm Fusion-RP 80 Å column (150 × 4.6 mm) was used for all chromatographic separations. Metabolite acquisition was done using Analyst Software, and. quantification and peak analysis were carried out using MultiQuant software (version 3.0.1). Complete metabolomic data from this study are provided in the Supplementary Worksheet 1.

### Statistical Analysis

Details of the statistical analysis employed to determine the significance of the data are provided in all the figure legends. All the data in the manuscript are represented as MeanSEM unless mentioned otherwise. For statistical analysis of the data, average mean fluorescence intensities of all the controls were compared to the test samples to compute the relative level of expression. N represents the number of biological replicates, and n represents the number of cells analysed in all the biological replicates of a particular condition. *P < 0.05, **P < 0.01, ***P < 0.001 and ****P < 0.0001.

## Acknowledgements

We acknowledge the use of all instruments at the mass spectrometry and confocal imaging (CIFF) facilities of NCBS (TIFR) and BRIC inStem. We thank members of the SL lab and Ramray Bhat for discussions and suggestions, and Vikas Trivedi for critical comments on the manuscript. We thank Drs. Srikala Raghavan, Minhaj Sirajuddin, Dhandapani Perundurai, and Dasaradhi Palakodeti for sharing resources. S.L. acknowledges funding support for this study from the DBT–Welcome Trust India Alliance (IA/S/21/2/505922), and the S. Ramachandran National Bioscience Award for Career Development from the Department of Biotechnology, Government of India.

## Author contributions

Conceptualization, U.A. and S.L.; methodology, U.A., J.S., S.A., and S.L.; investigation, U.A., J.S., S.A.; writing – original draft, U.A. and S.L.; writing – review and editing, U.A., J.S., S.A. and S.L.; funding acquisition, resources, and supervision, S.L.

## Conflict of interest

The authors declare no conflicts of interest.

---

**Supplementary Figure 1 (Related to Figure 1):**
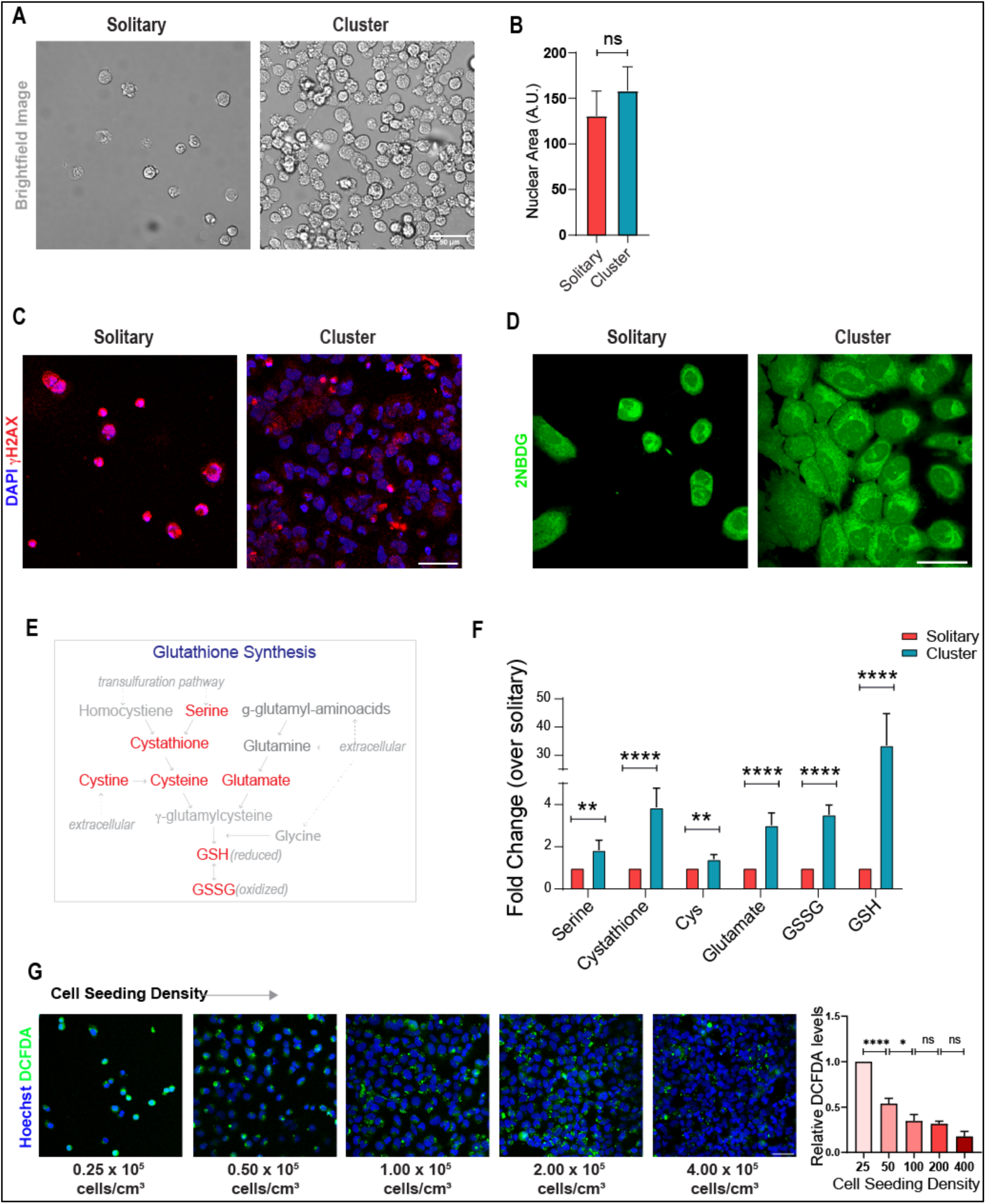
Metabolic and cell state of solitary and clustered cells grown on Poly-L-lysine-treated tissue culture dishes. A. Cell clustering at high seeding density: Bright field image showing clustering of cells when seeded at high density. B. Nuclear size comparison in solitary and clustered cells: Bar plot showing the mean nuclear area of solitary and clustered cells (N=3, Solitary (n) = 286, Cluster (n) = 988). C. Increased DNA damage marker in solitary cells: Immunocytochemistry image showing the expression of γH2AX in solitary and clustered cells. D. Glucose uptake in solitary and clustered cells: Immunocytochemistry image showing 2-NBDG intensities in solitary and clustered cells. No significant difference was observed between the two conditions. E. Pathway of glutathione synthesis – highlighting critical amino acids involved in red. F. Relative amounts of key amino acids and metabolites from the glutathione synthesis pathway, in solitary vs clustered cells. (N=3) G. DCFDA fluorescence in cells, as seeded at increasing cell densities (as indicated). The relative DCFDA fluorescence (normalized to the number of cells) is shown in the right bar graphs. (N=3) \**p* ≤ 0.05, \*\**p* ≤ 0.01, \*\*\*\**p* ≤ 0.0001, ns=not significant (student’s t test). All graphs are presented as mean ± SEM.

**Supplementary Figure 2 (Related to Figure 2):**
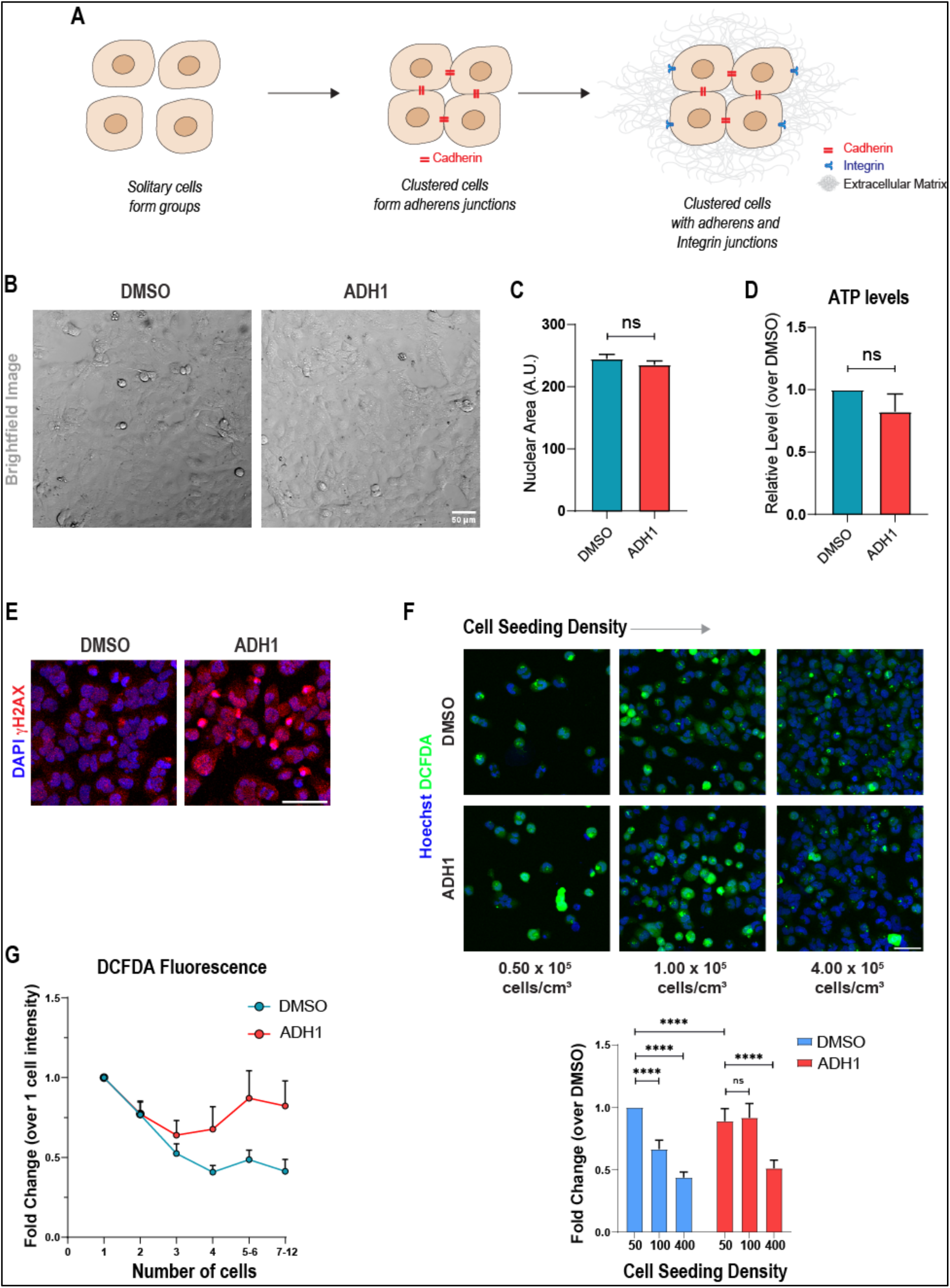
Metabolic and cell state of clustered cells treated with cadherin inhibitor ADH1 and DMSO control grown on Poly-L-lysine-treated tissue culture dishes. A. Sequential formation of cell–cell and cell–ECM junctions during clustering: Schematic showing that the formation of adherens junctions (cadherin-based) is one of the initial events stabilizing cell clusters, preceding the formation of cell–ECM junctions (integrin-based). B. Effect of ADH1 treatment on cell morphology: Bright field image showing clustered U2OS cells seeded in the presence of ADH1 and DMSO control. C. Effect of ADH1 treatment on cell size: Bar plot quantifying nuclear area—a proxy for cell size—in clustered cells treated with ADH1 and DMSO control (DMSO (n) = 972, ADH1 (n) = 1411). D. Effect of cell–cell junction perturbation on cellular ATP levels: Bar plot showing relative ATP levels in clustered cells upon ADH1 treatment, suggesting that disruption of cell–cell junctions does not cause immediate changes in cellular bioenergetics. (N=3). E. Effect of ADH1 treatment on DNA damage marker γH2AX: Immunocytochemistry image showing γH2AX expression in clustered cells treated with ADH1 compared to DMSO control, indicating elevated DNA damage upon disruption of cell–cell junctions (N=3). F. Effect of ADH1 treatment on ROS levels across densities: Immunocytochemistry images and corresponding bar plot showing that ADH1 treatment disrupts the density-dependent decrease in ROS levels, except at very high seeding densities where ROS levels remain low, likely due to additional effects of close cell clustering. N=3, DMSO 0.5x10^5^ (n) = 945, DMSO 1x10^5^ (n) = 2107, DMSO 4x10^5^ (n) = 2772, ADH1 0.5x10^5^ (n) = 831, ADH1 1x10^5^ (n) = 1635, ADH1 4x10^5^ (n) = 2596). G. ADH1 treatment prevents density-dependent ROS reduction: Graph showing intracellular ROS levels using DCFDA staining in ADH1 and DMSO control-treated cells with increasing cell cluster size. The changes are represented as fold changes over the DCFDA intensity of a single cell. N=3, DMSO (1 cell) n = 107, DMSO (2 cell) n = 80, DMSO (3 cell) n = 31, DMSO (4 cell) n = 23, DMSO (5-6 cell) n = 24, DMSO (7-12 cell) n = 11, ADH1 (1 cell) n = 73, ADH1 (2 cell) n = 67, ADH1 (3 cell) n = 30, ADH1 (4 cell) n = 19, ADH1 (5-6 cell) n = 2, ADH1 (7-12 cell) n = 7). \*\*\*\**p* ≤ 0.0001, ns=not significant (student’s t test). All graphs are presented as mean ± SEM.

**Supplementary Figure 3 (Related to Figure 2 and 3):**
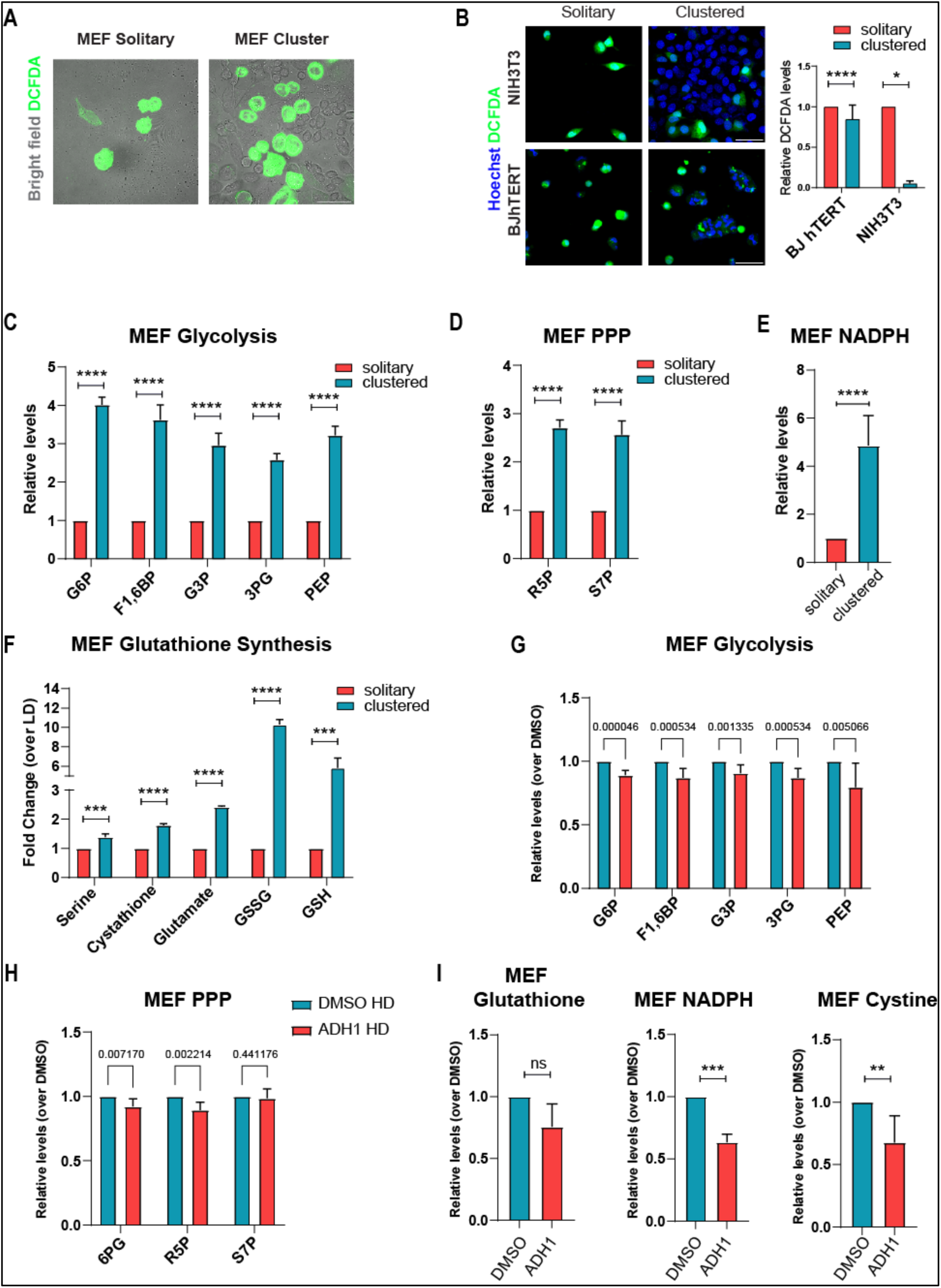
Metabolic and cell state of solitary and clustered MEFs, BJhTERT and NIH3T3 cells. (A) Cell clustering leads to a reduction in cellular ROS levels in MEF cells: Bright field image showing intensity of DCFDA staining. (B) Reduced ROS levels in clustered cells across additional cell lines (NIH3T3 and BJhTERT) : Immunocytochemistry images and corresponding bar plot showing DCFDA staining intensity in clustered cells, confirming a consistent reduction in intracellular ROS levels across multiple cell lines. (N=3, NIH3T3 Solitary (n) = 698, NIH3T3 Cluster (n) = 4454, BJhTERT Solitary (n) = 272, BJhTERT Cluster (n) = 938). (C) Clustering increases upper glycolysis metabolites in MEFs: Bar plot showing steady-state levels of upper glycolysis metabolites in clustered mouse embryonic fibroblasts (MEFs) compared to solitary MEFs. N=3. (D) Clustering increases PPP metabolites in MEFs: Bar plot showing levels of pentose phosphate pathway (PPP) metabolites in clustered MEFs relative to solitary MEF. N=3. (E) Clustering increases NADPH levels in MEFs: Bar plot showing NADPH levels in clustered MEFs, indicating enhanced reductive metabolism upon clustering. N=3. (F) Clustering promotes glutathione synthesis in MEFs: Bar plot showing levels of metabolites associated with glutathione synthesis in clustered MEFs. N=3. (G) ADH1 reduces upper glycolysis metabolites in clustered MEFs: Bar plot showing upper glycolysis metabolite levels in clustered MEFs treated with ADH1. N=3. (H) ADH1 reduces PPP metabolites in clustered MEFs: Bar plot showing PPP metabolite levels upon ADH1 treatment in clustered MEFs. N=3. (I) ADH1 reduces NADPH and glutathione pathway metabolites in clustered MEFs: Bar plot showing levels of NADPH and metabolites involved in glutathione and cystine metabolism in ADH1-treated clustered MEFs, highlighting the role of adherens junctions in supporting reductive metabolism. N=3. \**p* ≤ 0.05, \*\**p* ≤ 0.01, \*\*\**p* ≤ 0.001, \*\*\*\**p* ≤ 0.0001, ns=not significant (student’s t test). All graphs are presented as mean ± SEM.

**Supplementary Figure 4 (Related to Figure 4):**
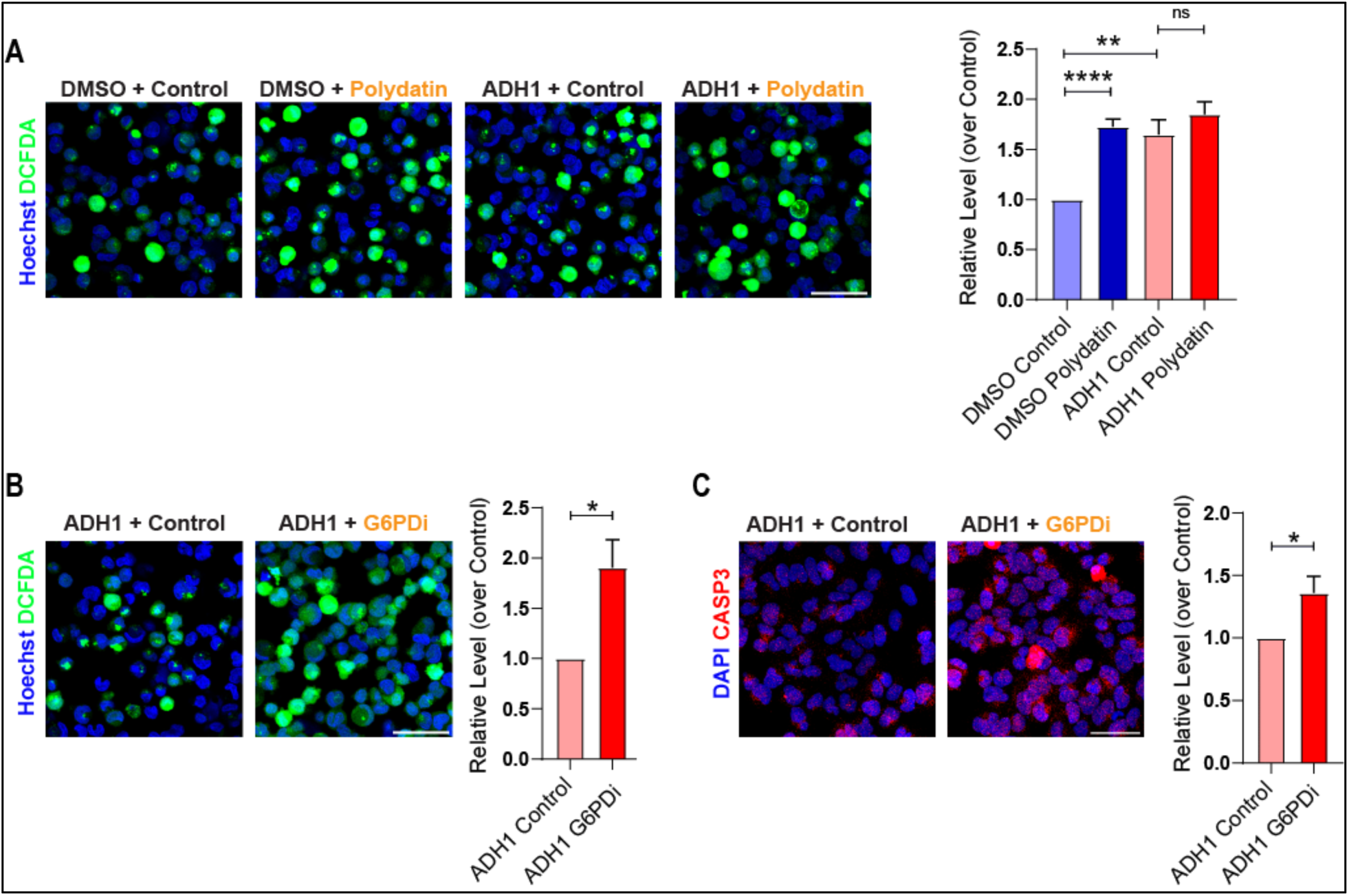
ROS expression and cell states of clustered cells treated with PPP and cadherin inhibitors. (A) Immunohistochemistry image showing ROS expression by DCFDA staining in clustered cells on inhibition of PPP by polydatin. (N=3, DMSO Control (n): 2834, DMSO Polydatin (n): 3893, ADH1 Control (n): 3951, ADH1 Polydatin (n): 3961). (B) Combined inhibition of PPP and adherens junctions increases ROS: Immunohistochemistry image and corresponding bar plot showing ROS expression by DCFDA staining in ADH1 treated cells on inhibition of PPP by G6PDi-1. (N=3, ADH1 + Control (n) = 2617, ADH1 + G6PDi (n) = 3276). (C) Combined inhibition of PPP and adherens junctions increases cell death marks: Immunohistochemistry image and corresponding bar plot showing caspase-3 expression in ADH1 treated cells on inhibition of PPP by G6PDi-1. (N=3, ADH1 + Control (n) = 1481, ADH1 + G6PDi (n) = 1799). \**p* ≤ 0.05, \*\**p* ≤ 0.01, \*\*\*\**p* ≤ 0.0001, ns=not significant (student’s t test). All graphs are presented as mean ± SEM.

**Supplementary Figure 5 (Related to Figure 6):**
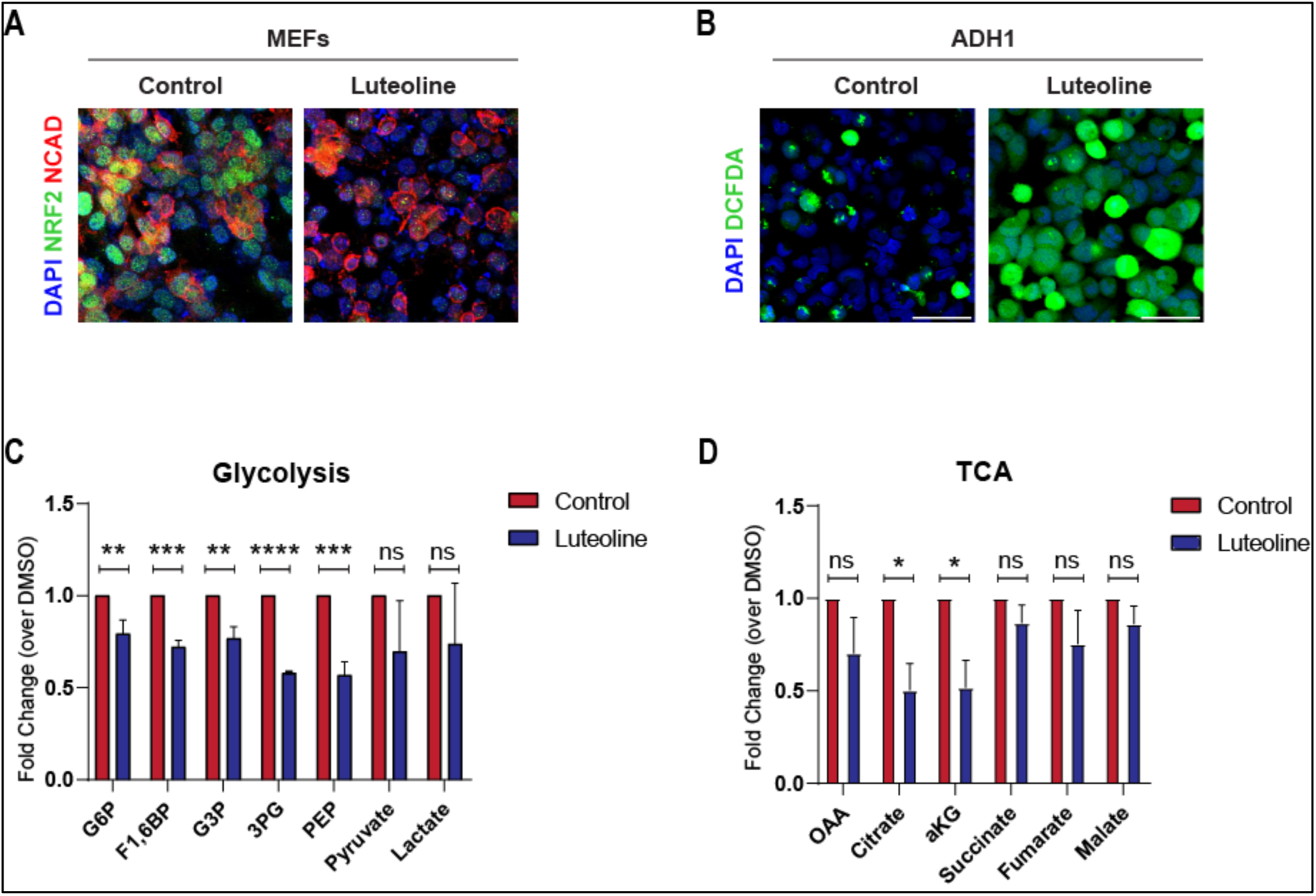
Metabolic state of clustered MEFs treated with cadherin junction inhibitor ADH1. (A) Luteoline treatment reduces NRF2 expression: Immunocytochemistry image showing decreased expression of NRF2 in cells treated with luteolin. (N=3) (B) Luteoline increases ROS in ADH1-treated clustered cells: Immunocytochemistry image showing elevated intracellular ROS levels in clustered cells treated with both luteoline and ADH1 by DCFDA staining. (N=3) (C) NRF2 inhibition reduces PPP metabolites: Bar plot showing levels of pentose phosphate pathway (PPP) intermediates in cells treated with luteoline. (N=3) (D) NRF2 inhibition reduces TCA metabolites: Bar plot showing levels of TCA metabolites in cells treated with luteoline. (N=3). \**p* ≤ 0.05, \*\**p* ≤ 0.01, \*\*\**p* ≤ 0.001, \*\*\*\**p* ≤ 0.0001, ns=not significant (student’s t test). All graphs are presented as mean ± SEM.

**Supplementary Table 1:** List of inhibitors used in the study.

| S.No. | Inhibitor | Source | Catalogue No. | Concentration |
| --- | --- | --- | --- | --- |
| 1 | ADH1-trifluoroacetate | Medchem<br>Express | HY-13541A | 0.1mM |
| 2 | Erastin | Medchem<br>Express | HY-15763 | 5mM |
| 3 | 2 deoxy-D-glucose | Sigma | D8375 | 0.5mM |
| 4 | Polydatin | Medchem<br>Express | HY-N0120A | 40mM |
| 5 | G6PDi-1 | Medchem<br>Express | HY-W107464 | 10mM |
| 6 | Luteolin | Medchem<br>Express | HY-N0162 | 0.1mM |

**Supplementary Table 2:** List of antibodies used in the study.

| S.No. | Antibody | Source | Catalogue No. | Concentration |
| --- | --- | --- | --- | --- |
| 1 | Cleaved Caspase-3 | R&D Systems | MAB835 | IC - 1:200 |
| 2 | Phospho-Histone H3 | Thermo Fischer Scientific | PA5-17869 | IC - 1:200 |
| 3 | N-Cadherin | Cell Signaling Technology | 13116 | IC - 1:400 |
| 4 | N-Cadherin | Cell Signaling Technology<br>(gifted by Dr. Minhaj Sirajuddin Laboratory at InStem) |  | IC - 1:400 |
| 4 | NRF2 | Cell Signaling Technology | 12721 | IC - 1:400 |
| 6 | Histone H2AX (p-Ser139) | Novus Biologicals<br>(gifted by Dr. Arjun Guha Laboratory at InStem) | NB100-384 | IC – 1:10000 |

## Notes

### Competing Interest Statement

The authors have declared no competing interest.

